# The synergistic effect of circRNA methylation promotes pulmonary fibrosis

**DOI:** 10.1101/2021.08.05.455186

**Authors:** Sha Wang, Wei Luo, Jie Huang, Menglin Chen, Jiawei Ding, Yusi Cheng, Wei Zhang, Shencun Fang, Jing Wang, Jie Chao

## Abstract

**Rationale:** N6-Methyladenosine (m^6^A) is the most common type of RNA methylation modification, mainly occurring on mRNA. Whether m^6^A-modified circRNAs are involved in different settings of pulmonary fibrosis remains unclear.

**Methods and Results:** Using an m^6^A-circRNA epitranscriptomic chip, candidate circRNAs were selected, in which hsa_circ_0000672 and hsa_circ_0005654 were specifically involved in SiO_2_-induced pulmonary fibrosis by targeting the same protein, eIF4A3, indicating that the m^6^A modification of these two circRNAs has a synergistic effect on fibroblast dysfunction induced by SiO_2_. A mechanistic study revealed that the m^6^A modification of circRNAs was mainly mediated by the methyltransferase METTL3. Furthermore, METTL3 promoted the activation, migration and activity of pulmonary fibroblasts and participated in SiO_2_-induced pulmonary fibrosis via circRNA m^6^A modification.

**Conclusion:** m^6^A methylation of circRNAs mediates silica-induced fibrosis via synergistic effects, enriching the understanding of circRNAs and uncovering a potential new target to treat fibrosis-related diseases.

## Introduction

Silicosis, the most common type of pneumoconiosis, is a pulmonary fibrosis disease caused by the long-term inhalation of large amounts of free silica (Leung *et al*, 2012). Inhaled silica particles can reach and persist in the peripheral lung (Sato *et al*, 2018). After being engulfed by alveolar macrophages (AMs), SiO_2_ triggers a series of inflammatory reactions, followed by fibrosis (Hoy & Chambers, 2020). The greatest challenges in silicosis diagnosis and treatment include the lack of a clear diagnostic target in the early stage and lack of effective treatment measures in the late stage. Therefore, exploring the pathogenesis of silicosis and identifying early diagnostic targets and late treatment measures are critical.

CircRNAs are a class of closed circular noncoding RNAs formed after trans-splicing. Because of the lack of 5′ or 3′ ends, they are relatively stable and can generally resist exonuclease degradation (Salzman, 2016). In recent years, the study of RNA epigenetics has become a popular approach in various fields, and RNA N6-methyladenosine (m^6^A) modification, the most abundant RNA modification in eukaryotes, has become a new research hotspot. m^6^A modification is the most common posttranscriptional modification of RNA in the transcriptome (Zaccara *et al*, 2019). m^6^A modification involves the attachment of a methyl group to the 6th nitrogen atom of adenine. m^6^A modification is also a dynamic process, including methylation and demethylation. Similar to the DNA methylation process, RNA m^6^A modification is catalyzed by methyltransferases and demethylases. Methyltransferases (“writers”) mainly include METTL3, METTL14, WTAP, KIAA1429, RBM15 and ZC3H13, while demethylases (“erasers”) mainly include FTO and ALKBH5. Additionally, readers, including YTHDF1, YTHDF2, and YTHDF3, recognize m^6^A-modified RNA and then perform functions (Zhuang *et al*, 2019). Studies have shown that m^6^A modification can regulate the stability, transport, splicing and translation of RNA at the molecular level (Lee *et al*, 2020; Wang *et al*, 2020b), and circRNAs also have various biological functions and are involved in the pathogenesis of Alzheimer’s disease, rectal cancer, depression, ischemia/reperfusion injury and other diseases (Chen *et al*, 2020; Li *et al*, 2018b; Lukiw, 2013; Ma *et al*, 2018; Zhang *et al*, 2020; Zhou & Yu, 2017). Recent studies have shown that m^6^A-modified circRNAs can maintain resistance to sorafenib by regulating β-catenin signaling in hepatocellular carcinoma (Xu *et al*, 2020b). Although a relationship between m^6^A and some tumors has been found in recent years, the mechanism of m^6^A modification in pulmonary fibrosis is unclear.

In this study, we investigated the role of m^6^A modification in pulmonary fibrosis, demonstrated the role of m^6^A-modified circRNAs in silicosis fibrosis, revealed a new regulatory model of circRNAs in silicosis, provided evidence of the correlation between m^6^A-modified circRNAs and disease, and enriched theoretical research on circRNAs.

## Methods

### Reagents

SiO_2_ particles with diameters of approximately 1-5 μm were purchased from Sigma-Aldrich (USA). Primary antibodies against METTL3, METTL14, WTAP, ALKBH5, FTO, α-SMA and FN1 were purchased from Proteintech. Antibodies against COL1A2 and GAPDH (MB001, Mouse) were obtained from Bioworld, Inc. (Louis Park, MN, USA). Antibodies against eIF4A3 were obtained from Affinity. Antibodies against p-PYK2 were obtained from ABCAM.

### Measurement of total m^6^A

The m^6^A levels of total RNA were measured using an EpiQuik m^6^A RNA Methylation Quantification Kit (Colorimetric) (Epigentek, Farmingdale, NY) following the manufacturer’s protocol. Total RNA was isolated from HPF-a cells and lung tissue using TRIzol reagent. In total, 200 ng of RNA was required for each sample.

### Agarose gel electrophoresis (AGE)

After the 1% agarose gel solidified, the comb was pulled out vertically. The gel was then placed into an electrophoresis tank, and the samples were run to a suitable position and stopped. The samples were placed in the gel imager for viewing and image capture.

### m^6^A RNA immunoprecipitation (MeRIP)

m^6^A RNA immunoprecipitation was performed using the Magna RIP Kit (Santa Cruz) according to the manufacturer’s instructions. Briefly, the cells were collected, and the same volume of RIP lysate was added to lyse the cells. The cells were placed on ice for 5 minutes, and the cells were completely lysed overnight at −80°C. The supernatant was centrifuged the following day. Next, magnetic beads were prepared for immunoprecipitation. The anti-m^6^A antibody and mouse IgG were incubated with the magnetic beads in buffer at room temperature for 30 min, and the cell lysate supernatant was mixed with the magnetic bead-antibody complex and incubated overnight at 4°C. The circRNA containing the m^6^A site was eluted on the second day, and the immunoprecipitated RNA was purified and extracted using phenol-chloroform-isoamyl alcohol. The extracted RNA was measured using the Nanodrop system for reverse transcription, followed by real-time PCR.

### Arraystar m^6^A-circRNA Epitranscriptomic chip high-throughput assay

The mouse model of silicosis was established according to the above method. The lung tissues of mice in the control group and model group were collected, and a high-throughput Arraystar m^6^A-circRNA Epitranscriptomic chip was used by Shanghai Kangcheng Biotech to investigate the differential m^6^A modification of circRNA in the lung tissues of mice in the control and model groups. The “m^6^A methylation level” for a transcript was calculated as the percentage of modified RNA (%Modified) in all RNAs based on the IP (Cy5-labeled) and Sup (Cy3-labeled) normalized intensities. The “m^6^A quantity” was calculated for the m^6^A methylation amount of each transcript based on the IP (Cy5-labeled) normalized intensities.

### Statistical analysis

GraphPad Prism 8.0.1 was used for statistical analysis, and all the data were expressed as means ± SD. T-test was used for comparisons between two groups, two-way ANOVA was used for comparisons between multiple groups, and significance was considered at a P value < 0.05.

## Results

### m^6^A-modified circRNAs in the lung tissue of silicosis model mice

The m^6^A modification level in circRNAs was detected by constructing a mouse model of silicosis. First, an Arraystar m^6^A-circRNA Epitranscriptomic microarray was used to investigate the differences in m^6^A modification in circRNAs in the lung tissues of mice in the silicosis and control groups (Figure 1A). Differences were found in the relative level of m^6^A-modified circRNAs and the relative quantity of m^6^A-modified circRNAs in the lung tissues of mice in the control and silicosis groups (Figure 1B–E). Among them, 24 were upregulated and 674 were downregulated in the relative level of m^6^A-modified circRNAs, 132 were upregulated and 336 were downregulated in the relative quantity of m^6^A-modified circRNAs. These results indicated that the differences in m^6^A modification in circRNAs might be involved in the pathological process of the lung after exposure to SiO_2_.

**Figure 1.**
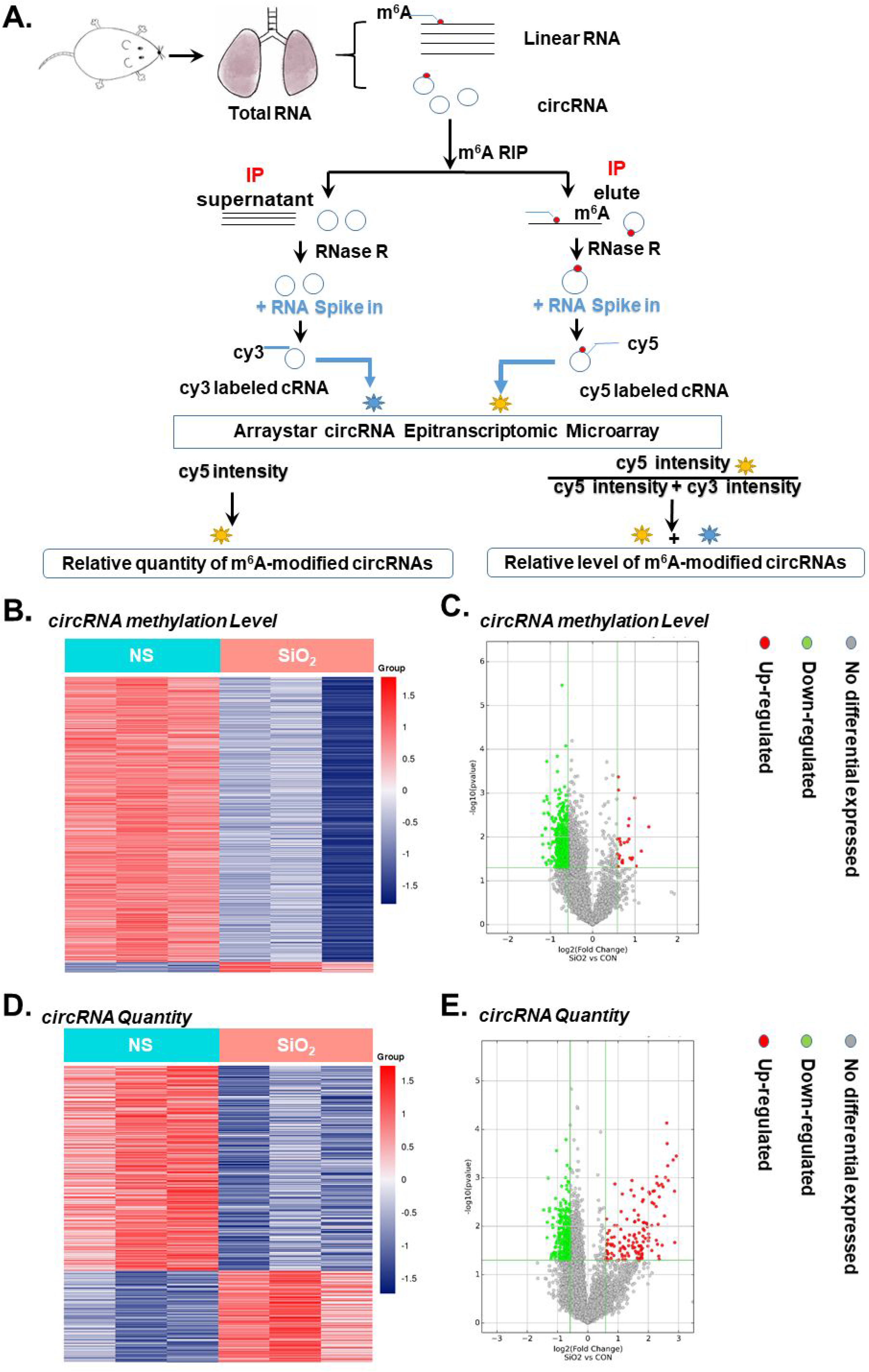
m^6^A modification of circRNAs in silicosis. **A** Arraystar m^6^A-circRNA epitranscriptomic microarray flow. **B** The heat map shows the difference in the level of m^6^A-modified circRNAs, and the red and blue strips indicate increased and decreased m^6^A levels, respectively. **C** The volcanic map shows the difference in the level of m^6^A-modified circRNAs. **D** Heat map showing the difference in the quantity of m^6^A-modified circRNAs. The red and blue strips indicate circRNAs with upregulated and downregulated expression, respectively. **E** The volcano map shows the difference in the quantity of m^6^A-modified circRNAs.

### Preliminary screening of circRNAs

To further understand the correlation of the m^6^A modification level between circRNA and total RNA, both the lung tissue of mice and human pulmonary fibroblasts treated with SiO_2_ were collected for total RNA m^6^A modification level detection. The m^6^A level in the lung tissue of the silicosis group was significantly higher than that in the control group by approximately 2 fold (Figure 2A), and the m^6^A modification level of total RNA in the human pulmonary fibroblasts treated with SiO_2_ increased sharply at 2 hours and then returned to normal levels (Figure 2B), indicating that m^6^A modification occurred in the pulmonary fibroblasts and might be involved in pulmonary fibrosis. Interestingly, the m^6^A modification level of total RNA in the plasma of healthy or silicosis patients showed no change (Figure E1), indicating that m^6^A modification occurred specifically in the lungs during pulmonary fibrosis. Having determined the upregulation of m^6^A modification, preliminary screening of circRNAs with increased m^6^A modification levels was conducted, in which 10 circRNAs homologous in humans and mice (circBase (http://www.circbase.org/) and circbank (http://www.circbank.cn/) databases; Table 1) with the most increased m^6^A modification levels were selected (Figure 2C). Next, the basic expression levels of the 10 circRNAs were detected, three of which had high expression levels (Figure 1C). The predicted m^6^A sites (SRAMP database; http://www.cuilab.cn/sramp) of the three circRNAs are shown. All 3 screened circRNAs had more than one m^6^A site. Among them, hsa_circ_0000672 and hsa_circ_0005654 had 9 and 7 m^6^A sites, respectively, while hsa_circ_0008336 had 2 m^6^A sites, hsa_circ_0000672 and hsa_circ_0005654 comprised 8 and 7 exons, respectively, and hsa_circ_0005654 comprised 4 exons (Figure 2D and Figure E2). These 3 candidate circRNAs were selected for further investigation.

**Figure 2.**
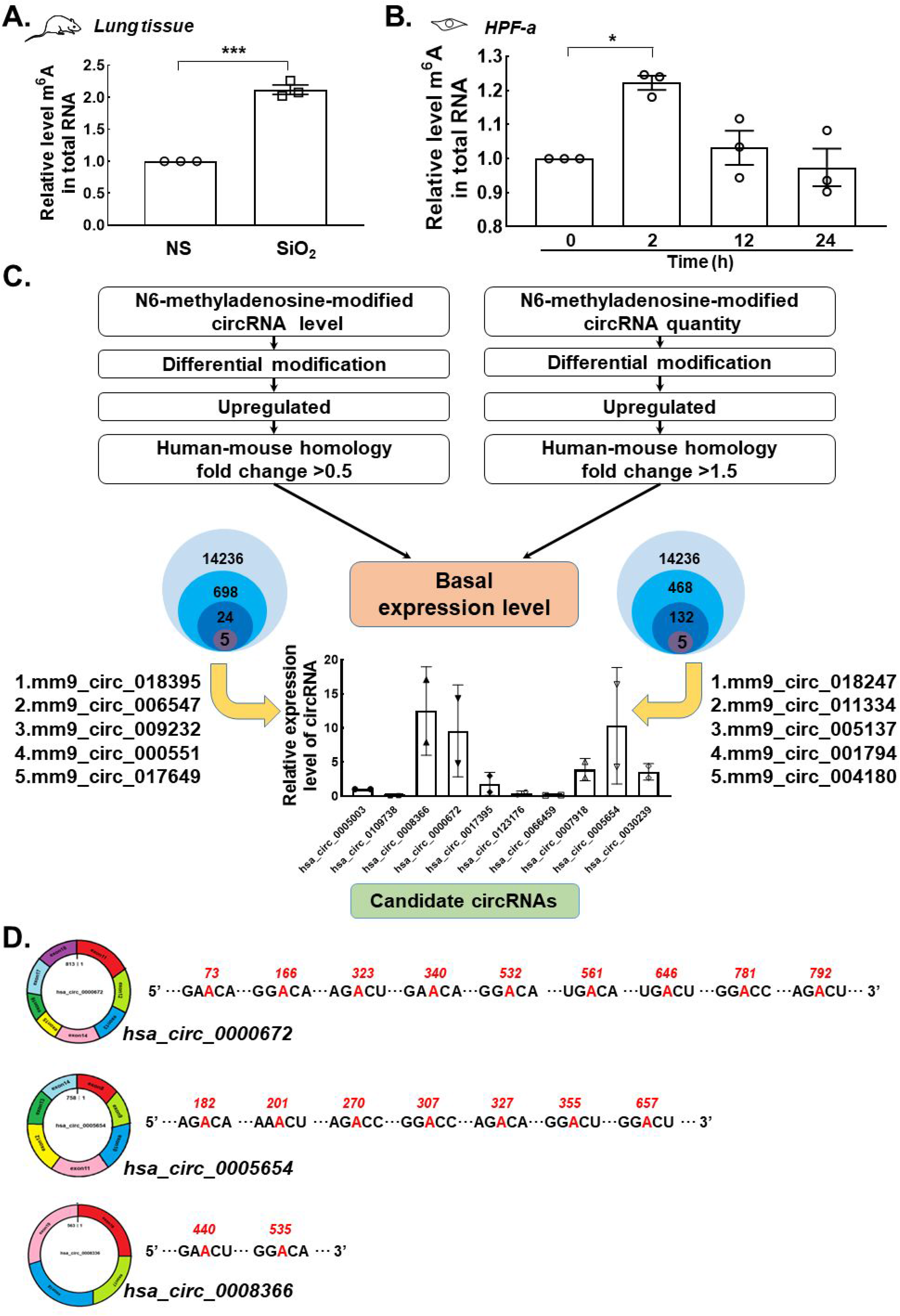
Preliminary screening of circRNAs. **A** The total RNA m^6^A level in the lung tissues of mice was detected using the EpiQuik m^6^A RNA Methylation Quantification Kit. **B** The EpiQuik m^6^A RNA Methylation Quantification Kit detects total RNA m^6^A levels in human pulmonary fibroblasts treated with SiO_2_ for different periods. **C** Screening process of circRNAs; the basic expression levels of 10 circRNAs in human pulmonary fibroblasts were detected by PCR. **D** Analysis of three circRNA m^6^A sites. The data are presented as means±SD. *P < 0.05, ** P < 0.01, *** P < 0.001.

**Table 1.**
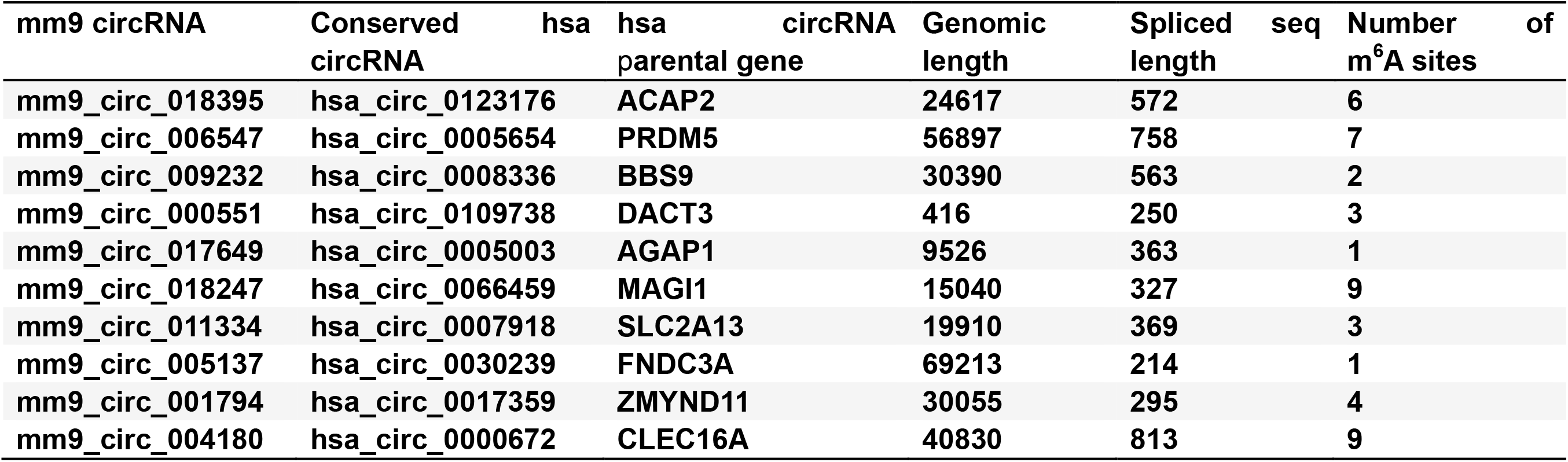
Candidate circRNAs homologous to humans and mice.

### hsa_circ_0000672 and hsa_circ_0000672 are involved in SiO_2_-induced fibroblast dysfunction

To determine the role of candidate circRNAs, siRNA was applied to knockdown their expression (Figure E2A). First, the effects of these three circRNAs on the activation of human pulmonary fibroblasts were detected, as indicated by changes in the fibroblast activation markers FN1, Collagen 1 and α-SMA. Unfortunately, the knockdown of hsa_circ_0000672, hsa_circ_0005654 (Figure E3B-E) and hsa_circ_0008336 (Figure E3F-I) alone did not affect the upregulation of FN1, Collagen 1 and α-SMA induced by SiO_2_. Furthermore, knockdown of these three circRNAs alone did not affect cell migration (Figure E4A and B). Considering the relatively low expression of most circRNAs (Li *et al*, 2018a), whether multiple circRNAs work together deserves to be investigated. Therefore, simultaneous knockdown with different combinations of candidate circRNAs was performed. Surprisingly, only hsa_circ_0000672 and hsa_circ_0005654 knockdown together abolished fibroblast activation induced by SiO_2_ (Figure 3A–D). Similarly, cell migration induced by SiO_2_ was significantly inhibited after the knockdown of these two circRNAs (Figure 3E and F). Taken together, the cooperation of hsa_circ_0000672 and hsa_circ_0005654 might be involved in SiO_2_-induced fibroblast dysfunction.

**Figure 3.**
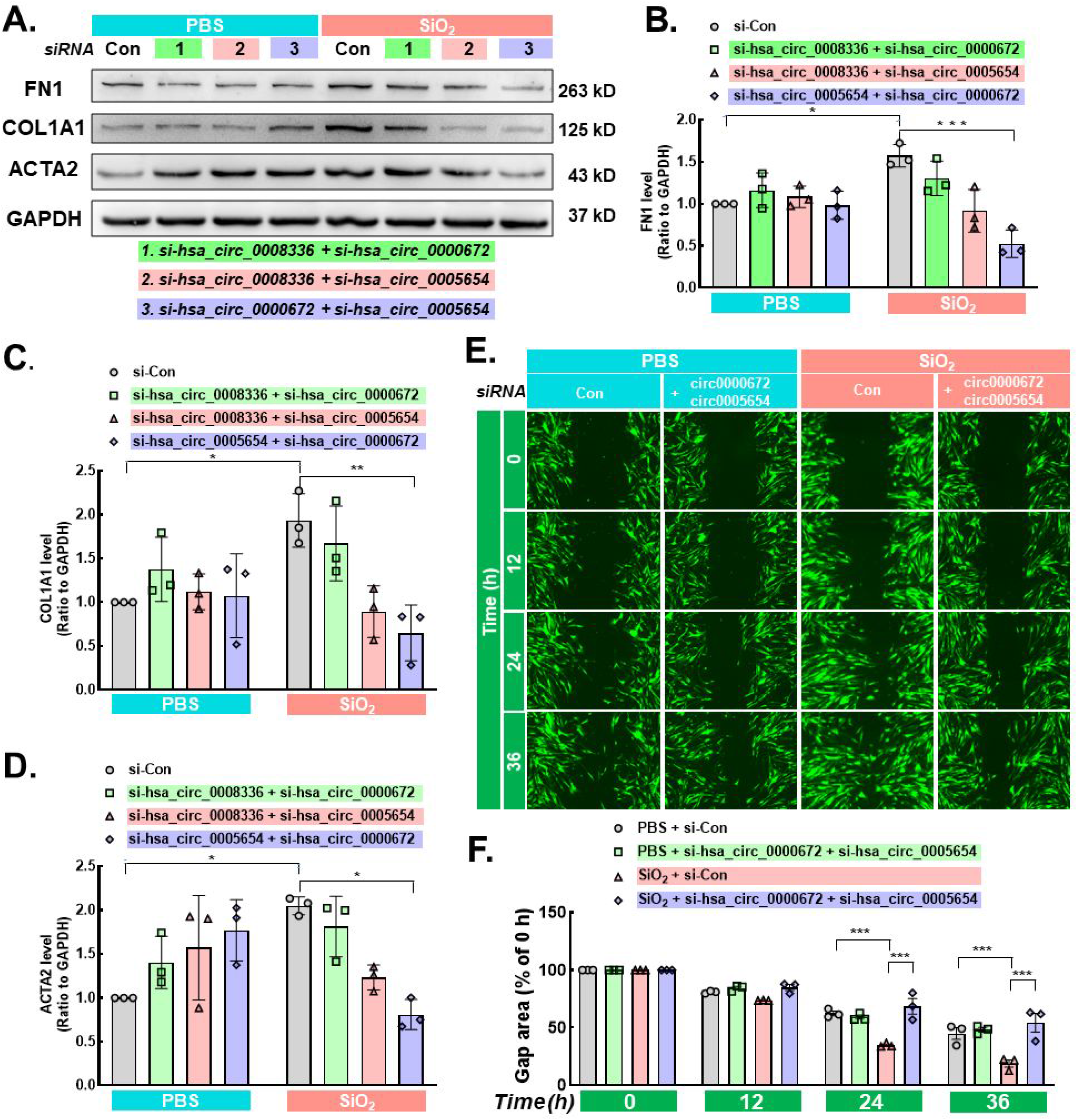
hsa_circ_0000672 and hsa_circ_0005654 promote the activation, migration and activity of human pulmonary fibroblasts. **A-D** WB was used to detect the effects of hsa_circ_0000672 and hsa_circ_0005654 knockdown on the human pulmonary fibroblast activation marker proteins FN1, Collagen 1 and α-SMA. **E-F** The effect of hsa_circ_0000672 and hsa_circ_0005654 knockdown on the migration of human pulmonary fibroblasts was examined using the wound healing assay. Scale=325 μm. The data are presented as means±SD. *P < 0.05, ** P < 0.01, *** P < 0.001.

### eIF4A3 is involved in SiO_2_-induced fibroblast dysfunction

We speculated why these two circRNAs are required for fibroblast dysfunction simultaneously. First, to confirm the existence of these two circRNAs, divergent and convergent primers to amplify hsa_circ_0000672 and hsa_circ_0005654 circular transcripts and linear transcripts were applied, respectively. PCR showed that hsa_circ_0000672 and hsa_circ_0005654 were detected only in cDNA but not in gDNA using divergent primers. Linear transcripts were amplified from both cDNA and gDNA using convergent primers (Figure E5A and B), indicating the existence of circRNAs in fibroblasts. As expected, SiO_2_ treatment did not change the expression of the two circRNAs (Figure 4A), indicating that epigenetic modification might be the main mode of action for these two circRNAs. Furthermore, in situ hybridization experiments demonstrated that hsa_circ_0000672 and hsa_circ_0005654 were mainly located in the cytoplasm of fibroblasts (Figure 4B). Next, to clarify why these two circRNAs show joint-work characteristics, circInteractome (https://circinteractome.nia.nih.gov/index.html) was applied to conduct bioinformatics analysis. A common RBP, eIF4A3, was revealed that could bind to the flanking intron sequences of these two circRNA pre-mRNAs (Figure 4C).

**Figure 4.**
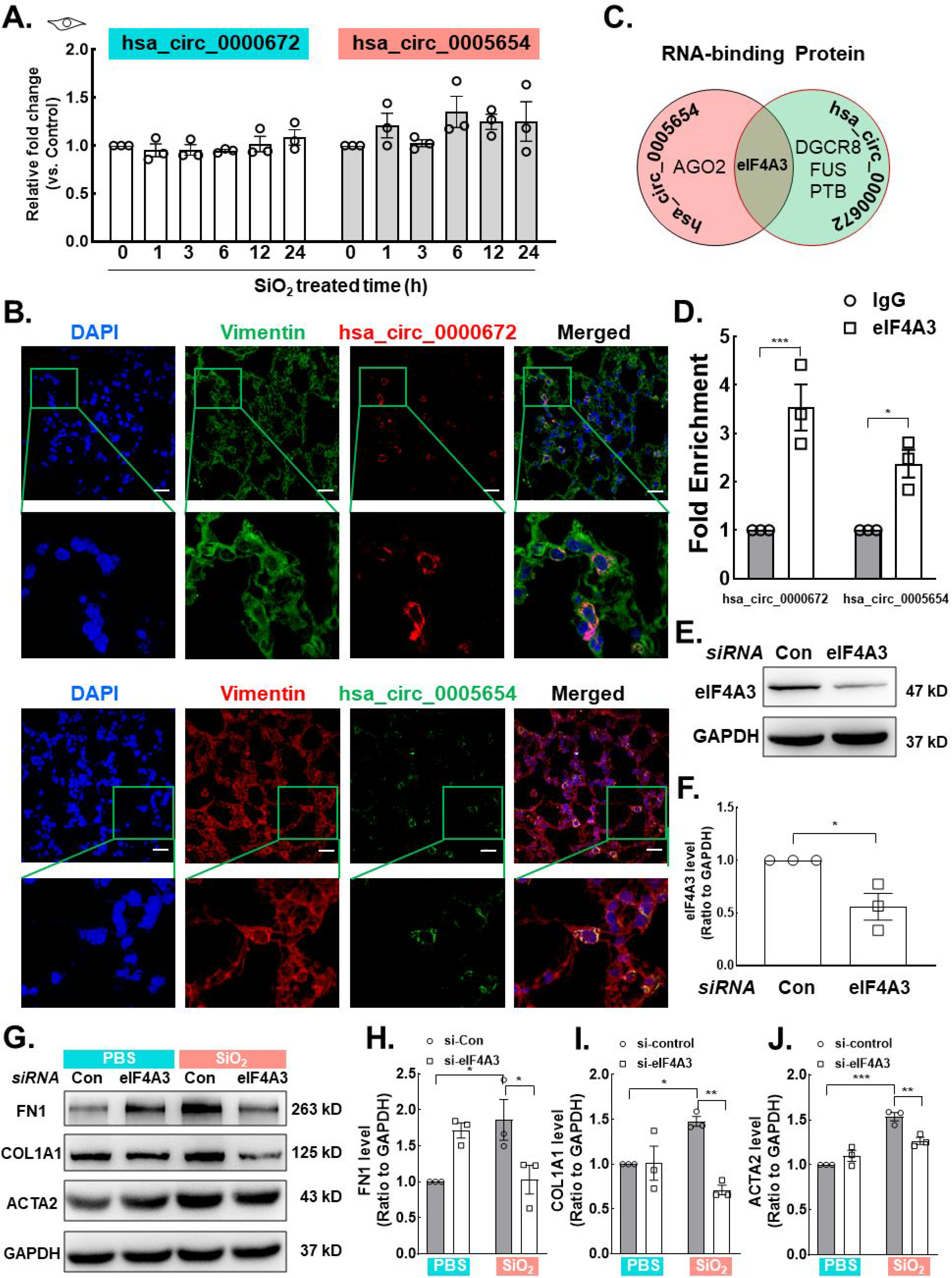
As the target proteins of hsa_circ_0000672 and hsa_circ_0005654, eIF4A3 promotes the activation, migration and activity of human lung fibroblasts. **A** PCR was used to detect the expression of hsa_circ_0000672 and hsa_circ_0005654 in human lung fibroblasts at different time points. **B** In situ hybridization showed that hsa_circ_0000672 and hsa_circ_0005654 were present in the cytoplasm. Scale=20 μm. **C** Bioinformatics analysis showed that hsa_circ_0000672 and hsa_circ_0005654 pre-mRNA have a common RBP. **D** The RIP test was used to detect the interaction of hsa_circ_0000672 and hsa_circ_0005654 with eIF4A3. **E-F** Knockdown efficiency of EIF4A3. **G-J** WB was used to detect the effects of knockdown of eIF4A3 on the human pulmonary fibroblast activation marker proteins FN1, Collagen 1 and α-SMA. The data are presented as means±SD. *P < 0.05, ** P < 0.01, *** P < 0.001.

eIF4A3 is a eukaryotic translation initiation factor that promotes the formation of circRNAs. Studies have found that eIF4A3 binds to MMP9 mRNA transcripts, inducing the cyclization of circMMP9 in GBM and increasing the expression of circMMP9 (Yue *et al*, 2019). However, previous results have suggested that the expression levels of hsa_circ_0000672 and hsa_circ_0005654 did not change with SiO_2_ exposure. Thus, the binding of eIF4A3 to the flanking intron sequence of circRNA pre-mRNA might not be a major factor. Therefore, we wondered whether these two circRNAs also interacted with eIF4A3. We verified through RIP experiments in 293T cells that hsa_circ_0000672 and hsa_circ_0005654 could bind to eIF4A3, confirming this hypothesis (Figure 4D). To further understand the role of eIF4A3 in fibroblast dysfunction induced by SiO_2_, knockdown of eIF4A3 was performed (Figure 4E and F), in which the upregulation of fibroblast markers after SiO_2_ exposure was attenuated (Figure 4G–J). The above results proved that hsa_circ_0000672 and hsa_circ_0005654 synergistically targeted eIF4A3, followed by fibroblast dysfunction.

### The level of METTL3 increases in vivo and in vitro

With the elucidation of the synergistic effect of two circRNAs, the upstream mechanism of m^6^A modification on circRNAs should be clarified. Because m^6^A modification involves methyltransferases and demethylases, which depend on the expression of m^6^A modification enzymes, the levels of five representative m^6^A modification enzymes were detected after SiO_2_ exposure in fibroblasts. Only the expression level of METTL3 showed an increase after SiO_2_ exposure (Figure 5A–B and Figure E6A-B), a finding that was confirmed by immunofluorescence staining (Figure 5C), while the other enzymes showed no change. Furthermore, the mRNA levels of the five enzymes were not affected by SiO_2_ exposure (Figure E6C), indicating that a nontranscriptional regulation mechanism might be involved. To validate the *in vitro* findings, immunofluorescence staining was conducted in the lung tissues of mice and human donors. The expression of METTL3 was elevated in the pulmonary fibroblasts of mice after seven and twenty-eight days of SiO_2_ exposure (Figure 5C). Consistent with these findings, immunofluorescence staining of lungs from healthy and silicosis patients also showed elevated METTL3 levels in pulmonary fibroblasts (Figure 5E). Taken together, these results indicated that METTL3 might be the main m^6^A modification enzyme in fibroblast dysfunction after SiO_2_ stimulation.

**Figure 5.**
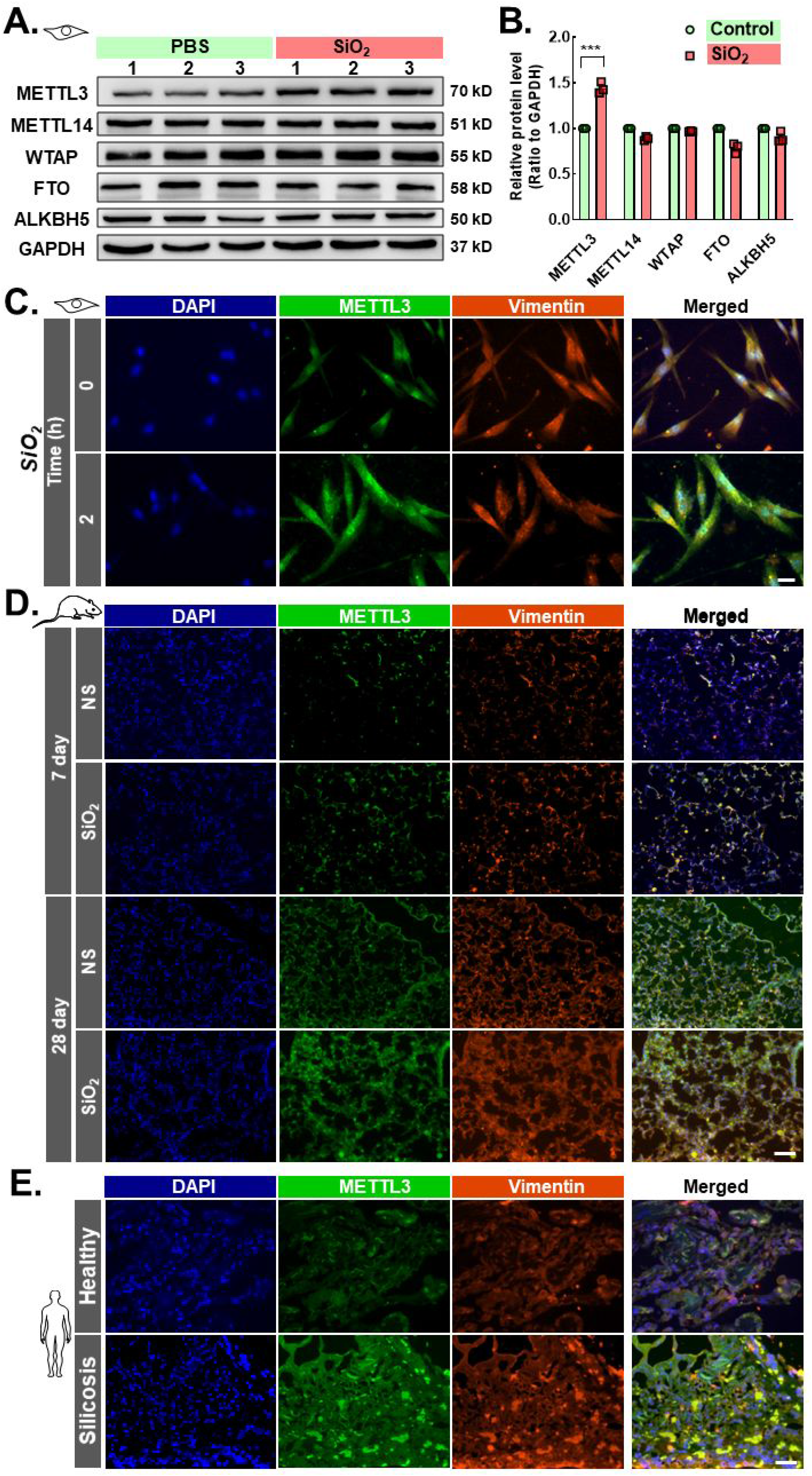
METTL3 expression is elevated in pulmonary fibrosis. **A-B** WB and q-PCR were used to detect changes in the five representative m^6^A modification enzymes in pulmonary fibroblasts. **C** The expression of METTL3 was detected by cellular immunofluorescence. Scale=30 μm. **D** The expression of METTL3 in the lung tissues of mice at 7 and 28 days after modeling was detected by immunofluorescence. Scale=50 μm. **E** The expression of METTL3 in the lung tissues of healthy individuals and silicosis patients was detected by immunofluorescence. Scale=50 μm. The data are presented as means±SD. *P < 0.05, ** P < 0.01, *** P < 0.001.

### METTL3 mediates RNA m^6^A modification and is involved in SiO_2_-induced fibroblast dysfunction

To further understand the effect of METTL3 on pulmonary fibroblasts, METTL3 siRNA in HPFs was applied (Figure E7A and B). Compared with the control group, METTL3 knockdown abolished the upregulation in the total RNA m^6^A level induced by SiO_2_ (Figure 6A), confirming the role of METTL3 in m^6^A modification. Furthermore, METTL3 knockdown significantly reduced the induction of fibroblast activation markers (Figure 6B–E) and fibroblast migration (Figure 6F and G). Additionally, phosphorylation of PYK2, a protein related to cell migration and invasion (Al-Juboori *et al*, 2019; Naser *et al*, 2018), was also attenuated after METTL3 knockdown (Figure E9A and B). Consistently, upregulation of cell viability induced by SiO_2_ was inhibited with METTL3 knockdown (Figure 6H). To further confirm the role of METTL3 in fibroblast activation, SAH (METTL3-METTL14 heterodimer complex inhibitor) (Li *et al*, 2016) was applied to pretreat fibroblasts for 2 hours. SAH abolished the increase in fibroblast activation markers induced by SiO_2_ (Figure 6I–L). Similarly, SAH pretreatment inhibited ascendant fibroblast viability induced by SiO_2_ (Figure 6M), a finding that was consistent with the result of METTL3 knockdown. Collectively, these experiments demonstrated that METTL3 mediates RNA m^6^A modification and is involved in SiO_2_-induced fibroblast dysfunction.

**Figure 6.**
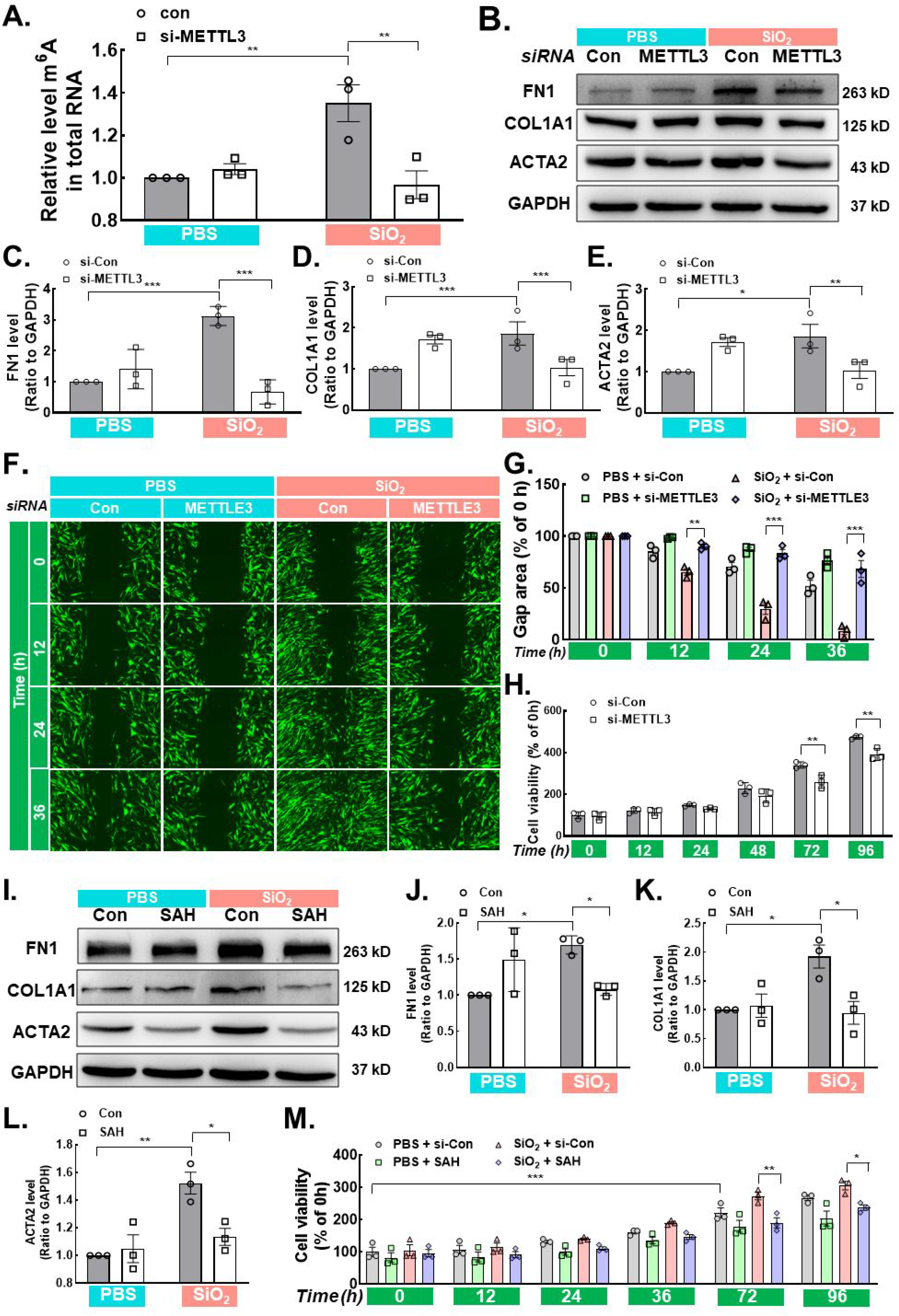
METTL3 mediates RNA m^6^A modification and promotes the activation, migration and activity of human pulmonary fibroblasts. **A** The EpiQuik m^6^A RNA Methylation Quantification Kit was used to detect total RNA m^6^A levels in human pulmonary fibroblasts after METTL3 knockdown. **B-E** WB was used to detect the effects of METTL3 knockdown on the human pulmonary fibroblast activation markers FN1, Collagen 1 and α-SMA. **F-G** The wound healing assay was used to detect the effect of METTL3 knockdown on the migration of human pulmonary fibroblasts. Scale=325 μm. **H** The CCK-8 assay was used to detect the effect of METTL3 knockdown on the activity of human pulmonary fibroblasts. **I-L** WB was used to detect the effects of the SAH inhibitor on the fibroblast activation markers FN1, Collagen 1 and α-SMA. **M** The CCK-8 assay was used to detect the effect of the SAH inhibitor on the viability of human pulmonary fibroblasts. The data are presented as means±SD. *P < 0.05, ** P < 0.01, *** P < 0.001.

### METTL3 mediates circRNA m^6^A modification, and m^6^A-modified circRNAs are involved in SiO_2_-induced fibroblast dysfunction

Having proven the role of circRNAs and METTL3 in fibroblast dysfunction, whether METTL3 mediates the m^6^A modification of circRNAs, as well as the m^6^A-modified circRNAs involved in SiO_2_-induced fibroblast dysfunction, must be clarified. *In situ* hybridization and immunofluorescence experiments indicated the colocalization of METTL3 with hsa_circ_0000672 and hsa_circ_0005654 (Figure 7A and Figure E8A and B). RIP experiments showed that METTL3 binds with hsa_circ_0000672 and hsa_circ_0005654 (Figure 7B). Furthermore, while METTL3 knockdown did not affect the expression of hsa_circ_0000672 and hsa_circ_0005654 (Figure 7C), METTL3 knockdown significantly reduced the m^6^A modification levels of circRNAs (Figure 7D), indicating that METTL3 was not involved in the expression of hsa_circ_0000672 and hsa_circ_0005654 but in the m^6^A modification of circRNAs.

**Figure 7.**
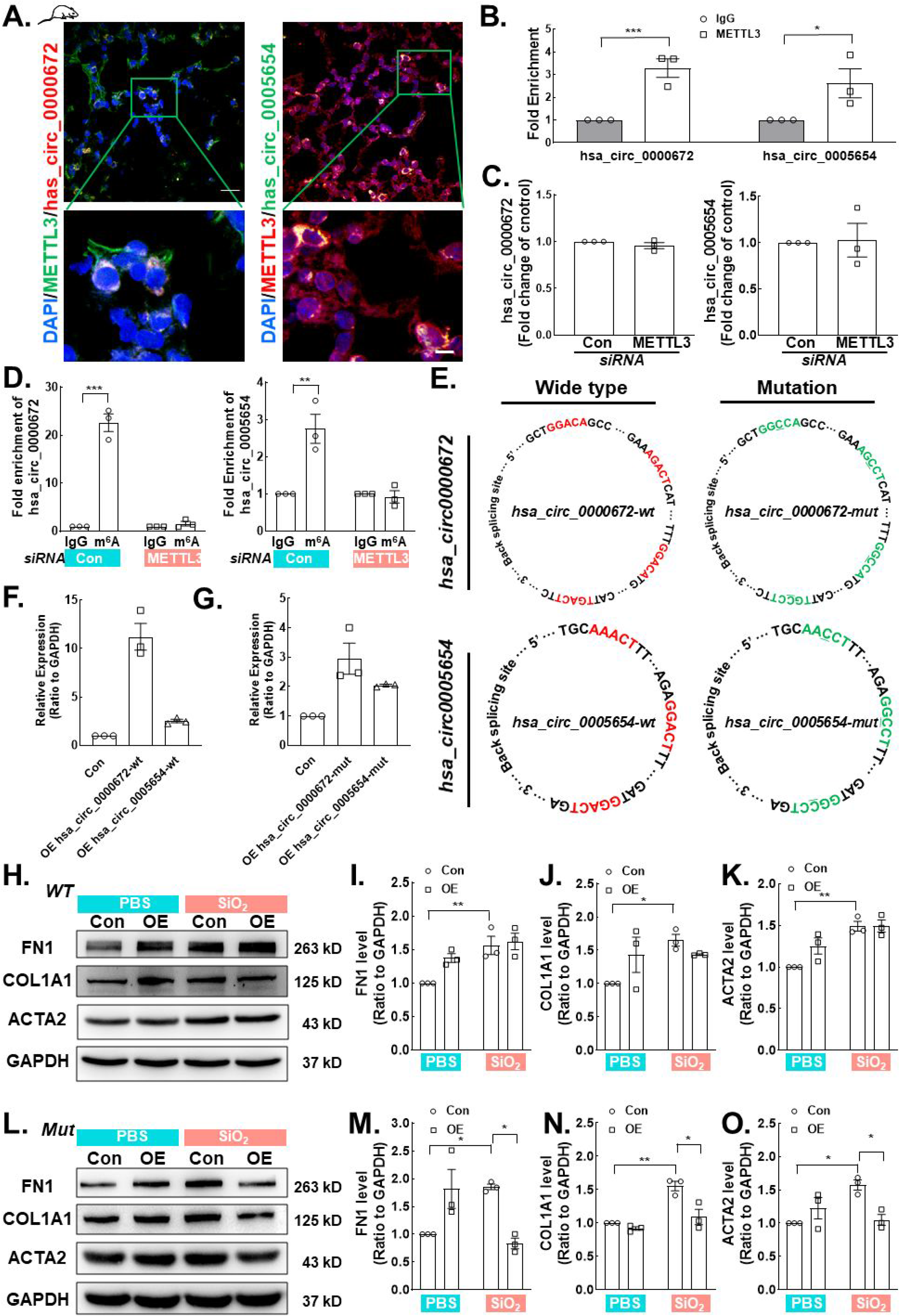
METTL3-mediated circRNA m^6^A modification promotes the activation of human pulmonary fibroblasts. **A** In situ hybridization detection of hsa_circ_0000672 and hsa_circ_0005654 colocalization with METTL3. Scale=20 μm. **B** RIP test of the interaction of METTL3 with hsa_circ_0000672 and hsa_circ_0005654. **C** Changes in circRNA m^6^A modification after METTL3 knockdown were detected by me-RIP. **D** PCR was used to detect the changes in hsa_circ_0000672 and hsa_circ_0005654 expression after knocking down METTL3. **E** Hsa_circ_0000672 and hsa_circ_0005654 with the m^6^A site mutated or wild-type overexpression plasmids were constructed. **F-G** The overexpression efficiency of wild-type and m^6^A site-mutated overexpression plasmids in human pulmonary fibroblasts was detected by PCR. **H-K** WB was conducted to detect the effect of the wild-type overexpression plasmid on the human pulmonary fibroblast activation markers FN1, Collagen 1 and α-SMA. **I-O** WB was conducted to detect the effect of the m^6^A site mutated overexpression plasmids on the human pulmonary fibroblast activation markers FN1, Collagen 1 and α-SMA. The data are presented as means±SD, *P < 0.05, ** P < 0.01, *** P < 0.001.

Furthermore, hsa_circ_0000672 and hsa_circ_0005654 with mutated m^6^A sites (Figure 7E) were applied to confirm the connection between m^6^A-modified circRNAs and fibroblast function. Hsa_circ_0000672 and hsa_circ_0005654 with the m^6^A site mutated or wild-type overexpression plasmids were constructed, respectively. Sequencing results proved that the overexpressed plasmid was successfully constructed and induced overexpression in HPF-a cells (Figure 7F and G) and 293T cells (Figure E11A and B).

We have previously shown that circRNA knockdown significantly reduced the increase in fibroblast activation markers after SiO_2_ exposure (Figure 3A–D). Here, to demonstrate that circRNA m^6^A modification mediates this inhibition, HPF-a cells were knocked down with hsa_circ_0000672 and hsa_circ_0005654, after which either wild-type or mutant plasmids were transfected into the cells. Transfection of the wild-type plasmid restored the effect of SiO2 on the FN1, Collagen 1, and α-SMA (Figure 7H–K). However, transfection of the mutant plasmid attenuated those effect induced by SiO_2_ (Figure 7L–O), confirming the role of the m^6^A-modified circRNAs in the regulation of SiO_2_-induced fibroblast dysfunction.

## Discussion

The role of circRNA in various diseases has been recognized, but its mechanism is more complex. The current study revealed a new pattern of circular RNA action-synergistic effects on circRNA methylation, providing new evidence to explain the precise regulatory mechanism of circular RNA in pulmonary fibrosis.

As widely expressed noncoding RNAs, circRNAs are produced by back-splicing. Because of their special circular structure, circRNAs are protected from exonuclease cleavage and degradation and thus have high stability (Kristensen *et al*, 2019). CircRNAs have various biological functions. They act as sponges for microRNAs, block mRNA translation, regulate transcription and selective splicing, interact with RBPs involved in translation. (Han *et al*, 2018) and participate in the pathogenesis of glioblastoma, hepatocellular carcinoma, breast cancer, colon cancer, lung cancer and other cancers (Han *et al.*, 2018; Lei *et al*, 2020; Zeng *et al*, 2018). CircRNA-002178 acts as a ceRNA to promote the expression of PDL1/PD1 in lung adenocarcinoma (Wang *et al*, 2020a). In gastric cancer, some oncogenic circRNAs promote cell proliferation, migration and invasion, and some anticancer circRNAs have also been found to be biomarkers for GC diagnosis and prognosis (Li *et al*, 2020). Additionally, circRNAs regulate energy metabolism in cancer, including glucose metabolism, lipid metabolism, amino acid metabolism, and oxidative respiration (Yu *et al*, 2019). The role of circRNA in pulmonary fibrosis was performed recently (Dai *et al*, 2021; Li *et al*, 2019). For example, circHECTD1 promotes pulmonary fibrosis via HECTD1-mediated pulmonary fibroblast activation (Chu *et al*, 2019) or ZC3H12A-induced macrophage activation (Zhou *et al*, 2018). Although most of these studies focused on the functional role of a certain circRNA, the current study reveals a novel mode of action of the circRNA-synergistic effect. Because the expression level of circRNA in cells is generally low (Chen *et al.*, 2020), the joint-work mode facilitates subtle control of circRNAs on downstream cascades. The mechanism of circRNA synergistic effects is complex. One explanation may be the circRNA composition because the clarified circRNAs in the current study have abundant m^6^A binding sites. Another explanation is the mechanism of action, such as sharing the same downstream target. In the current study, two circRNAs could bind the same protein, eIF4A3. Although two circRNAs have abundant m^6^A binding sites and the same target protein, cumulative synergistic effects, or “synergy of synergistic effects”, may account for the significant effect of the simultaneous knockdown of two circRNAs on fibroblast functions. Another reasonable assumption is that circRNAs with similar characteristics may not only have synergistic effects but also antagonistic or additive action, possibly explaining the subtle regulation of circRNAs.

Epigenetic modification is involved in various biological activities, including DNA methylation, RNA methylation, and histone modification. Recently, an increasing number of studies have been conducted on RNA methylation, in which N6-methyladenosine (m^6^A) is the most common type and has been reported to be involved in the occurrence and development of various diseases (Zeng *et al.*, 2018). In hepatoblastoma, m^6^A-modified mRNAs promote the proliferation of hepatoblastoma by regulating CTNNB1 (Liu *et al*, 2019). Recent studies have shown that m^6^A also occurs in long noncoding RNAs and circRNAs (Li *et al.*, 2020; Ni *et al*, 2019). m^6^A-modified circRNAs have cell type-specific expression, which is different from mRNA expression (Zhou *et al*, 2017). Additionally, m^6^A modification can regulate circRNA fate, such as regulating the translation of circRNAs, promoting the degradation of circRNAs and regulating the biogenesis of circRNAs (Chen *et al*, 2019; Li *et al.*, 2020; Yang *et al*, 2017). In the current study, high-throughput sequencing of an Arraystar m^6^A-circRNA epitranscriptomic microarray was applied to detect differences in circRNA m^6^A modifications. Interestingly, although the microarray results suggested a decrease in most circRNA m^6^A levels and quantities, the m^6^A level of total RNA showed a different pattern. Only a small proportion of circRNAs showed m^6^A modification patterns similar to those of the total RNA m^6^A modification. Because of the subtle regulation pattern of circRNAs, these circRNAs, consistent with total RNA, may exhibit a significant effect. Therefore, we selected circRNAs with elevated m^6^A modification levels for subsequent experiments.

The current study suggests that circRNA m^6^A modification is mainly mediated by METTL3, which may promote fibrosis via fibroblast activation. Previous studies have shown that METTL3-mediated m^6^A modification is crucial for epithelial-mesenchymal transformation and metastasis in gastric cancer (Yue *et al.*, 2019). METTL3-mediated m^6^A modification directly promotes YAP translation and increases YAP activity by regulating the MALAT1-miR-1914-3p-YAP axis, thereby inducing resistance and metastasis in NSCLC (Jin *et al*, 2019). Interestingly, we found that METTL3 is elevated at the protein level, but no change is found at the RNA level, indicating that METTL3 may also have modifications, such as posttranslational modifications and posttranscriptional modifications. METTL3 undergoes SUMOylation modification and promotes tumor progression by regulating Snail mRNA homeostasis in hepatocellular carcinoma (Xu *et al*, 2020a). The detailed mechanism of METTL3 in pulmonary fibrosis deserves further investigation.

Furthermore, METTL3-mediated m^6^A-modified hsa_circ_0000672 and hsa_circ_0005654 target eIF4A3, a eukaryotic translation initiation factor involved in circRNA biogenesis. For example, eIF4A3 binds to MMP9 mRNA transcripts, inducing the cyclization of circMMP9 in GBM and increasing the expression of cirMMP9, thereby promoting the proliferation, invasion and metastasis of GBM cells (Wang *et al*, 2018). Interestingly, eIF4A3 mediates the production of circSEPT9, which promotes cell growth and metastasis, thereby promoting the canceration and development of triple-negative breast cancer (Zheng *et al*, 2020). Additionally, hsa_circ_0030042 regulates abnormal autophagy by targeting eIF4A3 and protecting the stability of atherosclerotic plaques (Yu *et al*, 2021). These findings suggest the critical role of eIF4A3 in the biological function of circRNAs. In the current study, eIF4A3, an RBP, interacted with hsa_circ_0000672 and hsa_circ_0005654 to participate in pulmonary fibrosis. However, whether eIF4A3 affects the generation of other circRNAs in pulmonary fibrosis must be clarified.

In conclusion, SiO_2_ induces m^6^A modification of hsa_circ_0000672 and hsa_circ_0005654 via METTL3 in pulmonary fibroblasts; in turn, METTL3 targets eIF4A3 synergistically. eIF4A3 induces proliferation, migration and activation in pulmonary fibroblasts, followed by pulmonary fibrosis (Figure 8). The synergistic effects may be a considerable pattern of circRNA action, providing a new direction to explore the mechanism of pulmonary fibrosis.

**Figure 8.**
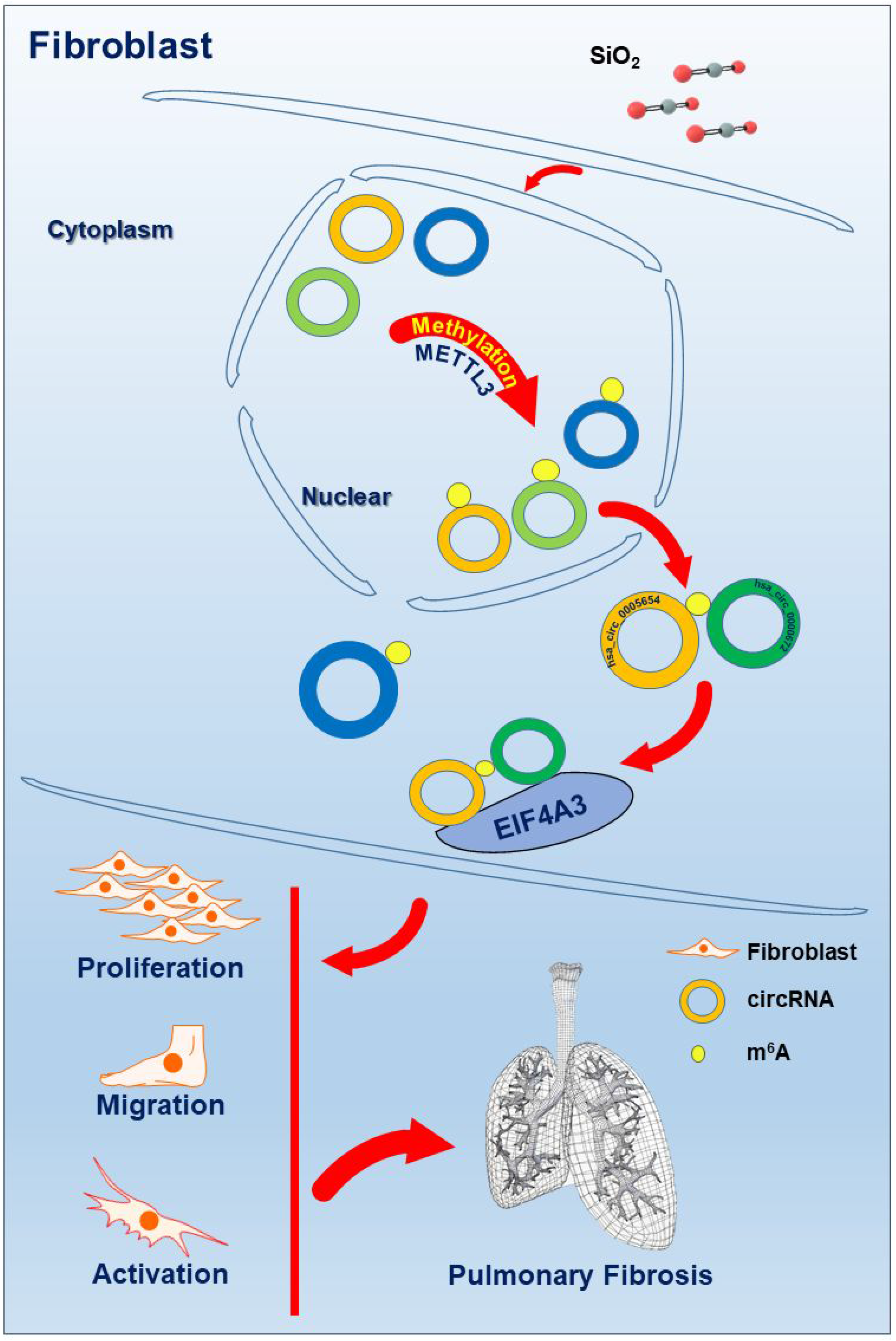
Schematic diagram showing the mechanisms through which silica induced m^6^A modification of hsa_circ_0000672 and hsa_circ_0005654, followed by activation of fibroblast.

## Acknowledgments

This study is the result of work that was partially supported by the resources and facilities at the core laboratory at the Medical School of Southeast University.

## Data sharing statement

All of the relevant raw data and materials are freely available to any scientist upon request.

